# Physical activity, aerobic fitness and brain white matter: their role for executive functions in adolescence

**DOI:** 10.1101/674846

**Authors:** Ilona Ruotsalainen, Tetiana Gorbach, Jaana Perkola, Ville Renvall, Heidi J. Syväoja, Tuija H. Tammelin, Juha Karvanen, Tiina Parviainen

## Abstract

Physical activity and exercise beneficially link to brain properties and cognitive functions in older adults, but it is unclear how these results generalise to other age groups. During adolescence, the brain undergoes significant changes, which are especially pronounced in white matter. Existing studies provide contradictory evidence regarding the influence of physical activity or aerobic-exercise on executive functions in youth. Little is also known about the link between both aerobic fitness and physical activity with white matter during puberty. For this reason, we investigated the connection between both aerobic fitness (20-m shuttle run) and physical activity (moderate-to-vigorous intensity physical activity) with white matter in 59 adolescents (12.7–16.2 years). We further determined whether white matter interacts with the connection of fitness or physical activity with three core executive functions (sustained attention, spatial working memory and response inhibition). Our results showed that only the level of aerobic fitness, but not of physical activity was related to white matter properties. Furthermore, the white matter of specific tracts also moderated the links of aerobic fitness and physical activity with working memory. Our results suggest that aerobic fitness and physical activity have an unequal contribution to the properties of white matter in adolescent brains. We propose that the differences in white matter properties could underlie the variations in the relationship between either physical activity or aerobic fitness with working memory.

**Highlights:**

- Aerobic fitness level, but not physical activity, is associated with white matter properties in several white matter tracts in the brain.
- The relationship between aerobic fitness and working memory was moderated by fractional anisotropy of the body of corpus callosum and in the right superior corona radiata.
- The relationship between physical activity and working memory was moderated by fractional anisotropy of the body and genu of corpus callosum.

## Introduction

Recent studies show that physical activity and high aerobic fitness beneficially link to many brain properties. Due to its significance for many cognitive functions, the brain’s white matter has been a target of increasing interest also in association with physical performance. A considerable amount of literature has been published about the connection between either physical activity or aerobic fitness with the brain’s white matter in older adults (e.g. Oberlin et al., 2016; Strömmer et al., 2018; Tian et al., 2015). However, investigative data concerning adolescents remain tentative. While the studies in older adults bring important practical implications, it is problematic to generalise these results to other age groups because certain white matter properties, such as fractional anisotropy (FA), which represents the degree of diffusion anisotropy and is used to assess microstructural changes in the brain white matter, change throughout an individual’s lifespan. During childhood and adolescence FA increases until it peaks at early adulthood followed by a gradual decline (Lebel et al., 2008; Westlye et al., 2010). Interestingly, it has been postulated that not only white matter properties (e.g. the FA) change with age but also its plasticity has been suggested to be different in younger and older individuals (Yotsumoto et al., 2014). This implies that our experiences and actions, such as physical activity, can have different impacts on our brains depending on age.

During puberty (10–17 years old), the body and brain undergo notable changes (Lebel et al., 2008; Spear, 2013). Due to differences caused by developmental changes, physical training may influence the brain differently depending on the childhood or adolescent phase. Earlier studies on young participants have focused on preadolescent children and older adolescents. However, to the best of our knowledge, no previous study has focused on the connection between either physical activity or aerobic fitness with white matter during puberty. In older adolescents, white matter FA, was not related to either aerobic fitness or self-reported physical activity (Herting et al., 2014). Yet, in the same study, the low physical activity group had fewer streamlines in the corticospinal tract and the forceps minor compared to the high physical activity group. On the contrary, studies focusing on preadolescent children have found associations between aerobic fitness, physical activity and white matter FA. In an exploratory study, Chaddock-Heyman et al. (2014) found that 9–10-year-old children with a higher fitness demonstrated a greater FA in the body of corpus callosum, superior corona radiata and superior longitudinal fasciculus. Two intervention studies demonstrated that physical activity intervention increased FA of the genu of corpus callosum in normal weight preadolescent children (Chaddock-Heyman et al., 2018) and of the uncinate fasciculus in overweight ones (Schaeffer et al., 2014) but not the FA of the superior longitudinal fasciculus (Krafft et al., 2014). Despite some inconsistencies in findings, these few studies suggest that besides the adult population, aerobic fitness and physical activity may be associated with white matter properties in young participants as well. However, care has to be taken when interpreting these early findings because the amount of studies involving child and adolescent participants remains scarce.

While physical activity and high aerobic fitness are reportedly beneficial for brain health, some studies further suggest exercise-related improvements in cognitive skills. Especially relevant in this context are executive functions: a set of processes highly important for cognitive control, and thus, also for domain-specific cognitive functions (for a review see Diamond [2013]). Several studies have suggested a relationship between either aerobic fitness or physical activity with executive function in youth (e.g. Chaddock et al., 2010; Davis et al., 2011; Hillman et al., 2014; Kamijo and Masaki, 2016; Subramanian et al., 2015). However, there are also studies that do not provide evidence for this association (e.g. de Greeff et al., 2016; Krafft et al., 2014; Schaeffer et al., 2014; Stroth et al., 2009; Tarp et al., 2016). It is currently unclear what might cause this variation between results. Interestingly, biological moderators have been suggested as a potential cause that affects the strength of the relationship between physical performance and cognition (Barha et al., 2017; Singh et al., 2018). Overall, the influence of physical activity and fitness on executive functions is likely to reflect complex pathways with several biological and psychological modulating factors. Identifying factors that moderate or alter the relationship between physical performance and executive functions will allow more individualised predictions and recommendations.

White matter integrity provides a factor possibly underlying the relationship between either physical activity or aerobic fitness with executive functions. The importance of white matter to cognition is widely known, and various studies report connections between white matter properties and the cognitive function throughout the lifespan (e.g. Chaddock-Heyman et al., 2013; Gold et al., 2010; Golestani et al., 2014; Mabbott et al., 2006; Nagy et al., 2004; Seghete et al., 2013). Additionally, studies investigating cognitive training suggest that white matter properties also predict the enhancement of cognitive skills after training and that the change in white matter properties following training relates to behavioural improvements (de Lange et al., 2017, 2016; Engvig et al., 2012; Mackey et al., 2012). Given the suggested relationship of also physical activity and fitness with white matter properties, it is conceivable that white matter has an important role in determining the manner in which physical performance associates with cognitive skills.

Previous studies do not provide direct answers to what extent exercise can influence brain white matter. For example, intervention studies suggest that exercise can increase white matter FA (Chaddock-Heyman et al., 2018; Krafft et al., 2014), but it may not increase equally at very high levels of exercise. Furthermore, this relationship between FA and exercise may further influence the cognitive benefits of physical exercise. Accordingly, we hypothesised that the relationship between either aerobic fitness or physical activity with core executive functions is different for individuals with low or high values of brain white matter FA. The main goals of the present study were (1) to examine if physical activity and aerobic fitness are related to white matter properties in 13−16-year-old adolescents and (2) to investigate whether white matter FA moderates the connection between either physical activity or aerobic fitness with core executive functions.

## Methods

### Participants

Participants (12.7–16.2 years old) for this cross-sectional study were recruited from a larger follow-up study (for more details, see Joensuu et al. [2018]). Sixty-one right-handed subjects participated in the brain magnetic resonance imaging (MRI) experiments, of which two participants did not complete the diffusion-weighted imaging protocol. A total of 59 subjects (39 female) were included in the analysis concerning physical activity, aerobic fitness, working memory, rapid visual information processing and white matter measures. From the 59 subjects, 58 were analysed for response inhibition as one participant did not complete the test. Participants were screened for exclusion criteria comprising MRI contraindications; neurological disorders; the use of medication that influences the central nervous system; any major medical condition; and left-handedness, which was assessed by the Edinburgh Handedness Inventory during the first research visit. Furthermore, to evaluate pubertal development, participants self-reported their stage of puberty by using the Tanner scale (Marshall and Tanner, 1970, 1969). The study was conducted according to the ethical principles stated in the Declaration of Helsinki, and prior to the participation in the study, each participant and his or her legal guardian provided written informed consent. The Central Finland Healthcare District Ethical Committee accepted the study. The participants were compensated with a 30-euro gift card for participating in the brain scans.

### Physical activity and aerobic fitness

The physical activity was objectively measured using the triaxial ActiGraph GT3X+ and wGT3X+ accelerometers (Pensacola, FL, USA; for full details, see Joensuu et al. [2018]). The participants were instructed to wear these devices on their right hip during waking hours for seven consecutive days (except during bathing and swimming). A valid measurement day consisted of at least 10 h of data. Subjects who had at least two valid weekdays and one valid weekend day were included in the analysis. For those subjects who did not meet these criteria, a multiple imputation method (explained in more detail below) was employed to compensate for the missing data. Activity counts were collected in 15-s epochs. When there was a period of at least 30 min of consecutive zero counts, it was considered as a non-wear period. Data were collected at a sampling frequency of 60 Hz and standardly filtered. A customised Visual Basic macro for Excel was used for data reduction. Cut points from Evenson et al.’s study were utilised in the analysis (Evenson et al., 2008; Trost et al., 2011). The moderate-to-vigorous intensity physical activity (MVPA) was converted into a weighted-mean value of MVPA per day ([average MVPA min / day of weekdays × 5 + average MVPA min / day of weekend × 2] / 7).

The maximal 20-m shuttle run test was employed to assess the aerobic fitness of the participants. The test was performed as described by Nupponen et al. (1999) and specified in detail for the present data collection in Joensuu et al. (2018). Each participant ran between two lines, 20 metres apart, at an accelerating pace, which was indicated with an audio signal. The duration that the participants ran until they failed to reach the end lines within two consecutive tones indicated their level of aerobic fitness. The speed in the first and second levels were 8.0 and 9.0 km/h, respectively. After the second level, the speed sequentially increased with 0.5 km/h per level. The duration of each level was one minute. The participants were verbally encouraged to keep running throughout the test.

In addition to the aerobic fitness test, the participants also completed a set of tests measuring muscular fitness (push-ups and curl-ups), flexibility and fundamental movement skills (a 5-leap test and a throwing-catching combination test), which are not included in the current study.

### Cognitive assessment

The participants completed a comprehensive cognitive test battery. In the present study, we focused on core executive functions because they have been associated with physical activity and fitness in previous studies that focused on different age groups. Thus, the participants in this study had to complete three different tests that measure core executive functions. Two of these tests were from the Cambridge Neuropsychological Automated Test Battery (CANTAB), including the Rapid Visual Information Processing (RVP) and Spatial Working Memory (SWM) tests (CANTAB eclipse version 6). The third test included a modified Eriksen Flanker task that measures response inhibition (Eriksen and Eriksen, 1974). Trained research assistants instructed the participants to perform the tablet-based tests according to the standard protocol in a silent environment.

For the RVP task of sustained attention, an array of numbers from 2 to 9 is presented in a pseudo-random order (100 digits per min). The participant’s task is to recognise three specific digit sequences (2-4-6, 3-5-7 and 4-6-8) and to press a response button when they detect the specific target sequence.

The SWM task assesses the participant’s ability to retain and manipulate visuospatial information. For this task, participants are asked to find a blue token hidden under a box, which is done by touching boxes on the screen. Once a blue token has been located, they are shown the same set of boxes and are asked to find the next one. The participants are also told that once a token has been found under a particular box that the same one would not hide any others tokens anymore. The difficulty of the task increases gradually from four to ten boxes. A principal component analysis was conducted for each CANTAB test, according to Rovio et al. (2016), to produce components for RVP and SWM that present the cognitive performance for each domain.

A modified Eriksen Flanker task was used to measure response inhibition. For this task, an array of five flanking fishes is shown to the participants, and they are asked to react as quickly and accurately as possible to the middle fish. We used four different conditions for this task: compatible congruent, compatible incongruent, incompatible congruent and incompatible incongruent. The first part of the test was compatible (congruent and incongruent), in which participants were asked to press the button at the side the fish was facing. For the congruent condition, all the other fishes in the array were swimming in the same direction as the middle fish, and for the incongruent condition, the other fishes were swimming in the opposite direction. The second part of the test was incompatible (congruent and incongruent), in which participants were asked to press the button at the opposite side of where the fish is facing. The average reaction time of the correct answers was used as an outcome measure in this task.

### Multiple imputation of the missing data

Multiple imputation was used to handle missing data. The proportion of missing values was 10% for the pubertal stage, 15% for the 20-m shuttle run and 22% for the MVPA. The reasons for missing values for most individuals included the absence from school during the measurement (e.g. due to sickness) and the insufficient number of valid measurement days (i.e. two weekdays and one weekend day) for physical activity. The incomplete data for several variables were imputed using the multiple imputation under a fully conditional specification (chained equations) (Van Buuren et al., 2006). The analysis was performed under the assumption of data missing at random as the crucial predictors – such as preceding measures (measured approximately six months before the current study) of pubertal stage, shuttle run tests (correlation with preceding 20-m shuttle run test = 0.57) and weekday measures of physical activity (correlation with the total MVPA [also weekend days included] = 0.95) – were available. As advised (Van Buuren, 2012 chapter 2.3.3), 50 imputed datasets were constructed and analysed. Each data set was constructed using 50 iterations of the multiple imputation by a chained equation algorithm to ensure the convergence of the iterative imputation process. The calculations were performed in R 3.4.0 (R Core Team, 2017) using the mice 2.3 package (Van Buuren and Groothuis-Oudshoorn, 2011). The model parameters and their standard errors were estimated for each imputed dataset and combined using Rubin’s rules (Van Buuren, 2012 p. 37–38) to obtain the final estimates of parameters and their standard errors. A full description of how the multiple imputation was done in the current study has been described previously in detail (Ruotsalainen et al., 2019).

### MRI acquisition

Imaging data were acquired on a 3T whole-body MRI scanner (MAGNETOM Skyra, Siemens Healthcare, Erlangen, Germany) using a 32-channel head coil at the Aalto NeuroImaging unit, Aalto University, Espoo, Finland. The total scanning time was approximately 45 min for the structural, diffusion-weighted, functional, field map and perfusion imaging. All scans, except perfusion MRI, were acquired using ‘Auto Align’ to minimise the variation in slice positioning (van der Kouwe et al., 2005). Prior to imaging the participants were familiarised with the measurement protocol. All participants were instructed to keep their head still during the scanning, and pads were used to minimise head motion. In addition, the participants wore earplugs to reduce the high noise caused by the MRI scanner.

For the diffusion-weighted imaging (DWI), we used a spin-echo based single-shot echo-planar (EPI) sequence with fat saturation. Prior to the DWI, high-order shimming was applied to reduce the inhomogeneities of the main magnetic field in the imaging area. In addition, the axial slices were slightly tilted in the anterior-posterior commissure line to avoid aliasing artefacts and artefacts caused by eye motion on the imaging slices. Seventy axial slices without gap were collected in 30 different diffusion gradient orientations. Two sets of diffusion-weighted images with b = 1000 s/mm2 and ten T2-weighted EPI images (b = 0 images) were acquired with two opposite phase encoding directions (anterior to posterior and posterior to anterior). The acquisition parameters were as follows: repetition time (TR) = 11,100 ms, echo time (TE) = 78 ms, field of view (FOV) = 212 mm, matrix size = 106, voxel size = 2 × 2 × 2 mm3, GRAPPA acceleration = 2 and phase partial Fourier = 6/8. The scanning time for each DWI series was 7 min 47 s.

### Image analysis

Diffusion-weighted images were processed using the FMRIB Software Library (FSL 5.0.11, www.fmrib.ox.ac.uk/fsl). For each subject, the magnetic susceptibility distortions were corrected using the topup tool (Andersson et al., 2003). Eddy current-induced distortions and subject movements were corrected using the eddy tool (eddy_cuda), including the slice-to-volume motion model and the outlier replacement (Andersson et al., 2017, 2016; Andersson and Sotiropoulos, 2016). This was followed by the removal of non-brain tissue using the Brain Extraction Tool (BET). Then, DTIFIT was used to fit the diffusion tensor model at each voxel. The output of the DTIFIT yielded voxel-wise maps of the FA, mean diffusivity (MD) and radial diffusivity (RD).

Voxel-wise statistical analysis of the FA, MD and RD data were carried out using Tract-Based Spatial Statistics (TBSS) – for methodological details, see Smith et al. (2006). First, every FA image was aligned to every other image to identify the ‘most representative’ image, which was used as a target image. Then, this target image was affine aligned into an Montreal Neurological Institute (MNI)152 standard space, and all subject’s FA data were transformed into MNI152 space by combining the nonlinear transform of the target FA image with the affine transform from that same target to the MNI152 space. Next, the mean FA image was created and thinned to create a mean FA skeleton that represents the centres of all tracts. Each subject’s aligned FA data was then projected onto this skeleton, and the resulting data fed into the voxel-wise cross-subject statistics. The skeleton was thresholded at an FA value of 0.2.

To test the relationship of physical behavioural measures (physical activity and aerobic fitness) with white matter tract measures(FA, MD and RD), we used FSL’s randomise tool with 10,000 permutations (Winkler et al., 2014). In this analysis, the age, pubertal stage and sex of the participant were used as covariates. The T-value difference in the voxel clusters was considered significant when the values passed – after the threshold-free cluster enhancement (TFCE) and family-wise error correction – a threshold of p < 0.05. The anatomical location of significant clusters was labelled with reference to the JHU ICBM-DTI-81 atlas. For the TBSS analysis, we used the average values of the imputed datasets (for the physical activity, aerobic fitness and pubertal stage). We also conducted a TBSS analysis using the head motion (eddy_restricted_movement_rms) as a covariate; however, this had negligible effects on the results. The analysis pipeline is available at Open Science Framework (https://osf.io/rg6zf/).

### Regression and moderation analyses

#### Aerobic fitness, physical activity and executive functions

Multiple linear regression, taking into account all 50 imputed datasets, was used to analyse the associations between core executive functions, physical activity and aerobic fitness. The following multiple regression model was used:

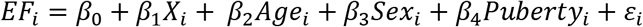

In the model, *EF*_*i*_ is the score of the executive function test of each test for a subject *i*, *X*_*i*_ is the physical activity or aerobic fitness, *Age*_*i*_, *Sex*_*i*_ and *Puberty*_*i*_ are the age, sex and pubertal stage, respectively and ε_i_ is the error term. The error variables were independent and identically distributed normal random variables with zero mean and the same standard deviation. All predictors were entered simultaneously into the model. The residual plots and Q-Q plots were used to check the assumptions of linearity as well as the normality and homoscedasticity of the residuals. In addition, homoscedasticity was tested using the Breusch–Pagan test (bptest) from the R package lmtest (Breusch and Pagan, 1979; R Core Team, 2017; Zeileis and Hothorn, 2002). The means of the residuals in all models were close to zero. The highest pairwise correlation was found between the pubertal stage and an age r of 0.59, indicating that the multicollinearity was not a factor of concern. All variance inflation factors were below a value of 2. We used the false discovery rate (FDR) to adjust for the multiple comparisons, and the results that had an alpha level smaller than 0.05 after the FDR adjustment were considered noteworthy (Benjamini and Hochberg, 1995).

#### Aerobic fitness, physical activity, FA and executive functions

A moderation analysis was applied to examine whether white matter FA in predetermined tracts changed the association of either aerobic fitness or physical activity with core executive functions. Based on previous literature investigating the relationship between either aerobic fitness or physical activity with white matter FA in young participants, we used the following white matter tracts as regions of interest: the body and genu of corpus callosum, the bilateral superior corona radiata, the bilateral superior longitudinal fasciculus and the bilateral uncinate fasciculus (Chaddock-Heyman et al., 2018, 2014; Schaeffer et al., 2014). These regions were masked from the white matter skeleton using the JHU ICBM-DTI-81 atlas, and the mean FA value was extracted for each region of interest. For the moderation analysis, we used the same model that was used for assessing the associations between core executive functions and either aerobic fitness or physical activity; however, an interaction term (FA*physical activity or FA*aerobic fitness) was added to the model. The variables involved with the interaction term were mean centred. When a significant interaction effect was detected, we conducted a simple slopes analysis to assess the relationships at high (+1 SD) or low (−1 SD) levels of the moderator. All statistical analyses were performed using the R 3.4.0 software (R Core Team, 2017) with the moderation analysis; the mitml 0.3-7 package was utilised (Grund et al., 2019). The results with an alpha level smaller than 0.05 were considered noteworthy.

## Results

### Participant demographics

Table 1 describes the participant demographics of the 59 participants (58 for the Flanker task).

**Table 1.**
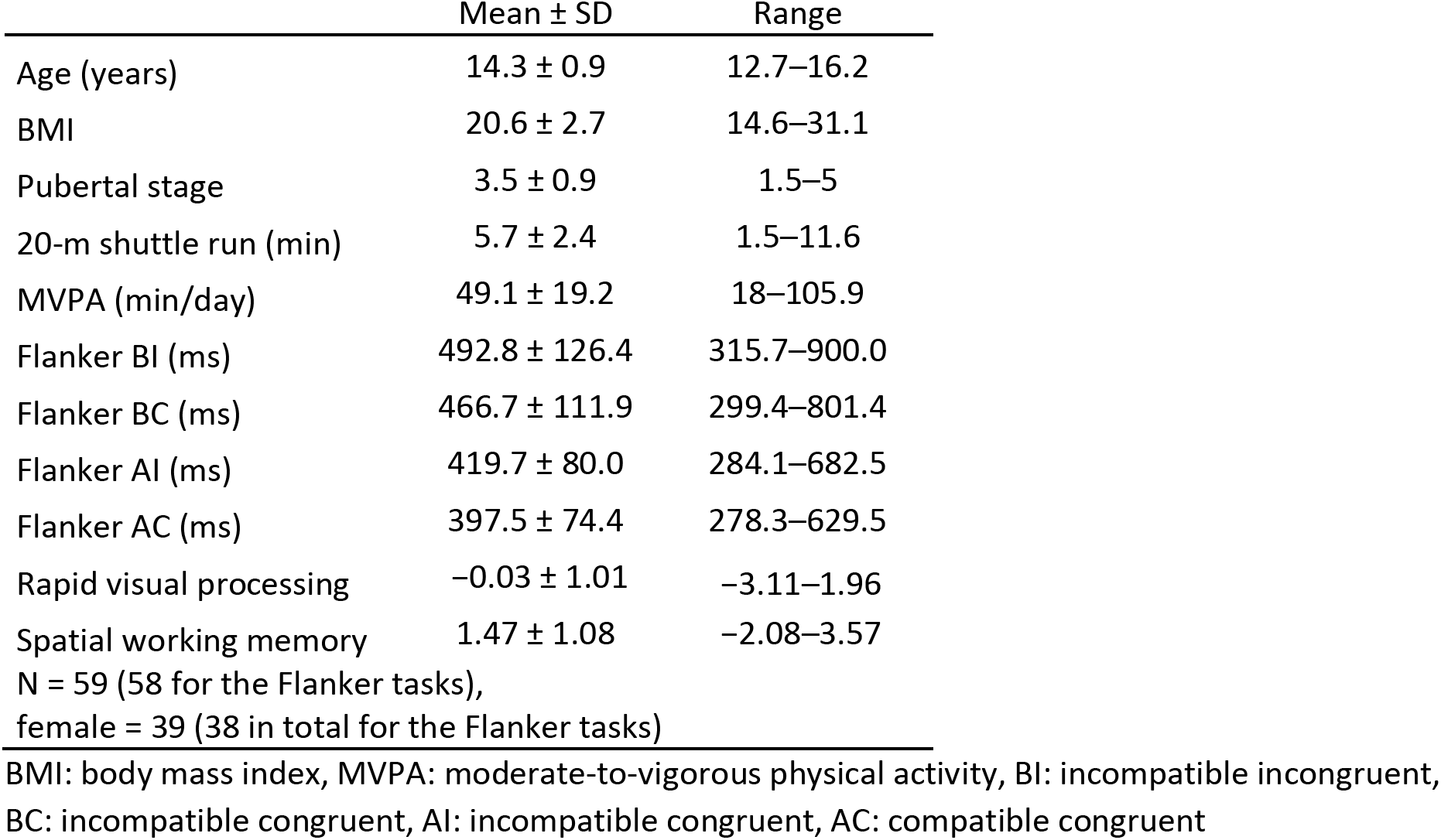
Participant demographics.

### Associations between white matter, aerobic fitness and physical activity

The TBSS analysis showed that aerobic fitness was positively associated with FA in several white matter tracts (Fig. 1 and Table 2). Aerobic fitness was associated with eight FA clusters of which the centres of mass localised to the left and right superior corona radiata and the body of corpus callosum. Besides FA, higher aerobic fitness was associated with a lower MD and RD in several white matter areas. These areas also included the corpus callosum and the bilateral superior corona radiata, although the associations were more widespread than those for FA. A complete list of clusters with MNI coordinates and overlapping anatomical tracts is given in Table 2. There were no associations between physical activity and FA, MD or RD. All statistical maps can be found in our Neurovault collection (https://neurovault.org/collections/5206/) (Gorgolewski et al., 2015).

**Fig 1.**
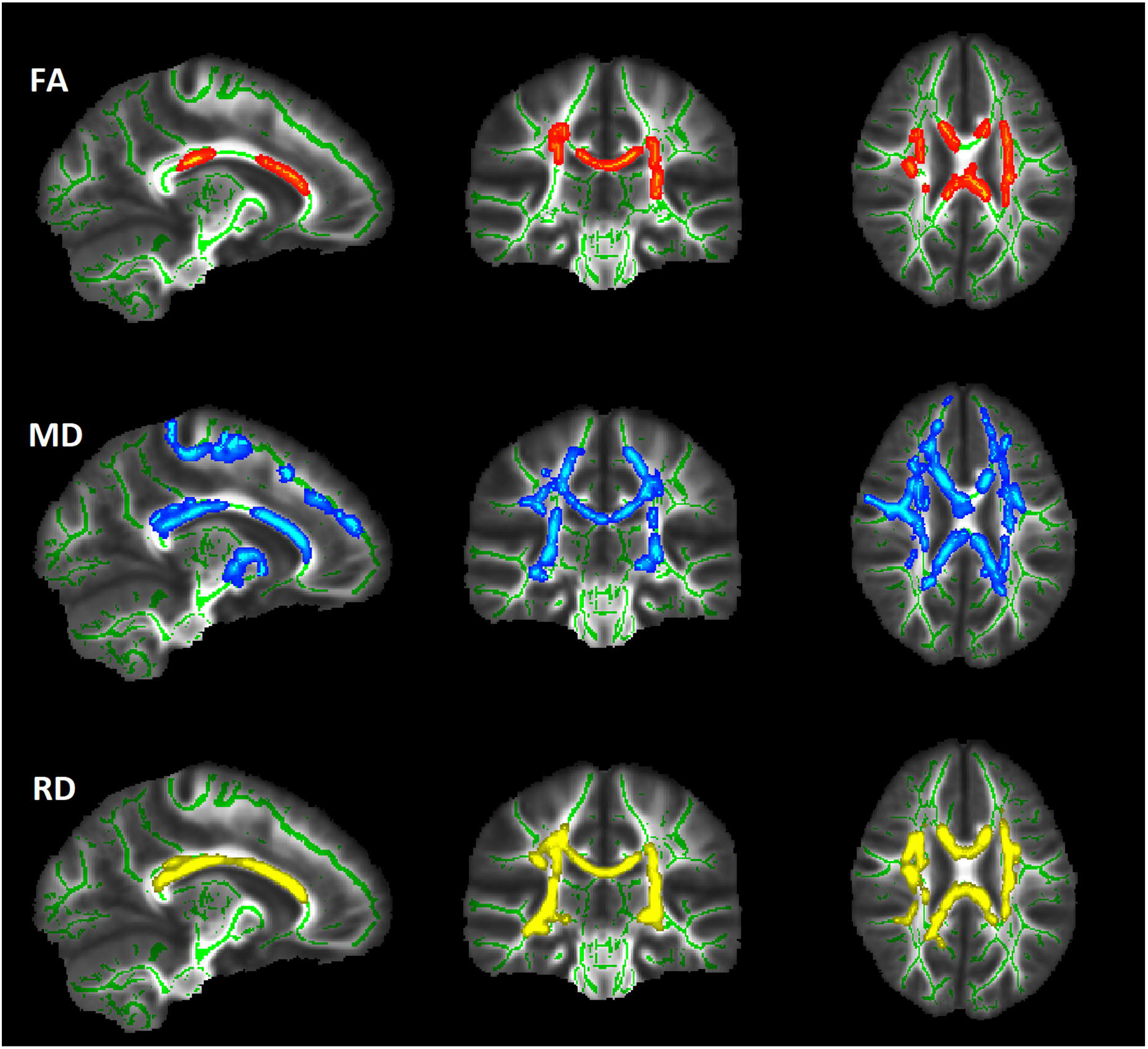
Associations (p < 0.05, corrected for family-wise error) between aerobic fitness and fractional anisotropy (FA), mean diffusivity (MD) and radial diffusivity (RD). The results are overlaid on an MNI152 1-mm template (MNI coordinates of the all slices are −12, −24 and 25). The association between aerobic fitness and FA (red) was positive, and the association between aerobic fitness, MD (blue) and RD (yellow) was negative. The significant regions are thickened for illustrative purposes.

**Table 2.**
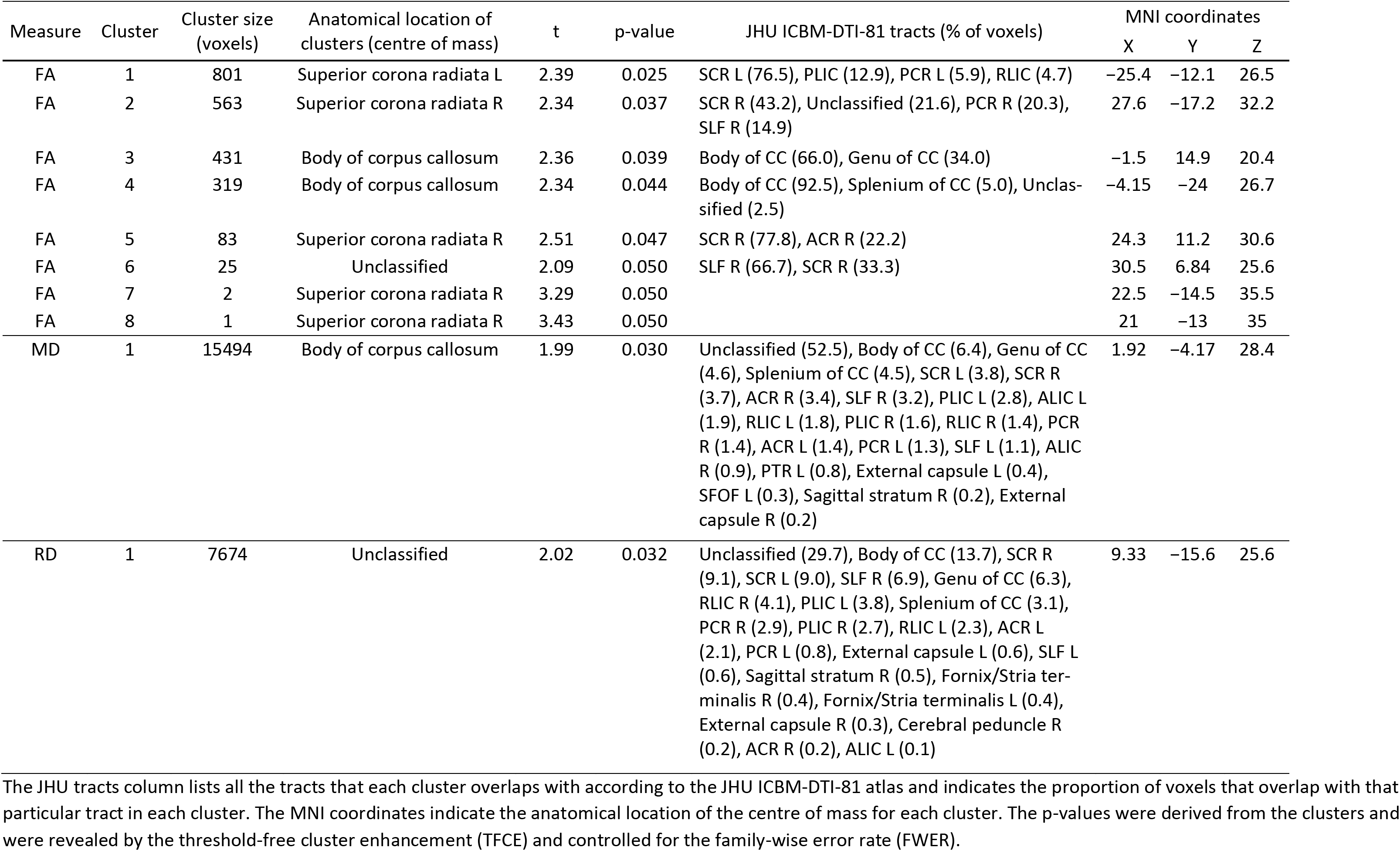
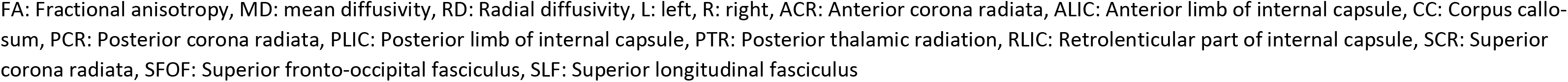
Characteristics of clusters that correlated with aerobic fitness.

| Measure | Cluster | Cluster size (voxels) | Anatomical location of clusters (centre of mass) | t | p-value | JHU ICBM-DTI-81 tracts (% of voxels) | MNI coordinates |  |  |
| --- | --- | --- | --- | --- | --- | --- | --- | --- | --- |
|  |  |  |  |  |  |  | X | Y | Z |
| FA | 1 | 801 | Superior corona radiata L | 2.39 | 0.025 | SCR L (76.5), PLIC (12.9), PCR L (5.9), RLIC (4.7) | -25.4 | -12.1 | 26.5 |
| FA | 2 | 563 | Superior corona radiata R | 2.34 | 0.037 | SCR R (43.2), Unclassified (21.6), PCR R (20.3), SLF R (14.9) | 27.6 | -17.2 | 32.2 |
| FA | 3 | 431 | Body of corpus callosum | 2.36 | 0.039 | Body of CC (66.0), Genu of CC (34.0) | -1.5 | 14.9 | 20.4 |
| FA | 4 | 319 | Body of corpus callosum | 2.34 | 0.044 | Body of CC (92.5), Splenium of CC (5.0), Unclassified (2.5) | -4.15 | -24 | 26.7 |
| FA | 5 | 83 | Superior corona radiata R | 2.51 | 0.047 | SCR R (77.8), ACR R (22.2) | 24.3 | 11.2 | 30.6 |
| FA | 6 | 25 | Unclassified | 2.09 | 0.050 | SLF R (66.7), SCR R (33.3) | 30.5 | 6.84 | 25.6 |
| FA | 7 | 2 | Superior corona radiata R | 3.29 | 0.050 |  | 22.5 | -14.5 | 35.5 |
| FA | 8 | 1 | Superior corona radiata R | 3.43 | 0.050 |  | 21 | -13 | 35 |
| MD | 1 | 15494 | Body of corpus callosum | 1.99 | 0.030 | Unclassified (52.5), Body of CC (6.4), Genu of CC (4.6), Splenium of CC (4.5), SCR L (3.8), SCR R (3.7), ACR R (3.4), SLF R (3.2), PLIC L (2.8), ALIC L (1.9), RLIC L (1.8), PLIC R (1.6), RLIC R (1.4), PCR R (1.4), ACR L (1.4), PCR L (1.3), SLF L (1.1), ALIC R (0.9), PTR L (0.8), External capsule L (0.4), SFOF L (0.3), Sagittal stratum R (0.2), External capsule R (0.2) | 1.92 | -4.17 | 28.4 |
| RD | 1 | 7674 | Unclassified | 2.02 | 0.032 | Unclassified (29.7), Body of CC (13.7), SCR R (9.1), SCR L (9.0), SLF R (6.9), Genu of CC (6.3), RLIC R (4.1), PLIC L (3.8), Splenium of CC (3.1), PCR R (2.9), PLIC R (2.7), RLIC L (2.3), ACR L (2.1), PCR L (0.8), External capsule L (0.6), SLF L (0.6), Sagittal stratum R (0.5), Fornix/Stria terminalis R (0.4), Fornix/Stria terminalis L (0.4), External capsule R (0.3), Cerebral peduncle R (0.2), ACR R (0.2), ALIC L (0.1) | 9.33 | -15.6 | 25.6 |

The JHU tracts column lists all the tracts that each cluster overlaps with according to the JHU ICBM-DTI-81 atlas and indicates the proportion of voxels that overlap with that particular tract in each cluster. The MNI coordinates indicate the anatomical location of the centre of mass for each cluster. The p-values were derived from the clusters and were revealed by the threshold-free cluster enhancement (TFCE) and controlled for the family-wise error rate (FWER).
FA: Fractional anisotropy, MD: mean diffusivity, RD: Radial diffusivity, L: left, R: right, ACR: Anterior corona radiata, ALIC: Anterior limb of internal capsule, CC: Corpus callosum, PCR: Posterior corona radiata, PLIC: Posterior limb of internal capsule, PTR: Posterior thalamic radiation, RLIC: Retrolenticular part of internal capsule, SCR: Superior corona radiata, SFOF: Superior fronto-occipital fasciculus, SLF: Superior longitudinal fasciculus

### Associations between physical activity, aerobic fitness and core executive functions

Multiple linear regression analysis revealed no significant associations between core executive functions (rapid visual information processing, spatial working memory and response inhibition) and either aerobic fitness or physical activity (Supplementary Table 1).

### White matter FA as a moderator of links between either aerobic fitness or physical activity with core executive functions

Finally, to test the hypothesis that white matter FA moderates the relationship between either physical activity or aerobic fitness with the core executive functions, we conducted an exploratory moderation analysis. We found that white matter FA in the body of corpus callosum and in the right superior corona radiata moderated the relationship between the 20-m shuttle run performance and spatial working memory. The interaction effect for FA of the body of corpus callosum*20-m shuttle run was β = −5.56, t = −2.64, p = 0.012, 95% confidence interval (CI) = −9.82, −1.30. For the FA of the right superior corona radiata*20-m shuttle run, this was β = −5.25, t = −2.14, p = 0.038, 95% CI = −10.19, −0.31. In addition, white matter FA in the body of corpus callosum and in the genu of corpus callosum moderated the association between the MVPA and spatial working memory. The interaction effect for FA of the body of corpus callosum*MVPA was β = −0.81, t = −2.52, p = 0.016, 95% CI = −1.47, −0.16. For the FA of the genu of corpus callosum, this was *MVPA β = −0.65, t = −2.30, p = 0.026, 95% CI = −1.23, −0.08.

A follow-up simple slopes analysis was performed to examine the nature of the interaction (Fig 2). These results revealed that the simple slopes were negative with high FA values and positive with low FA values. More specifically, with high FA values in the body of corpus callosum, the aerobic fitness was negatively associated with working memory (β = −0.21, t = −2.19, p = 0.034, 95% CI = −0.41, −0.02). Nevertheless, with low FA values, there were no significant associations between aerobic fitness and working memory. Aerobic fitness did not significantly associate with working memory, neither with high or low FA values in the right superior corona radiata. Regarding the relationship between physical activity and working memory, we found that with low FA values in the body and genu of corpus callosum, physical activity was positively related to working memory (low FA in the body of corpus callosum: β = 0.03, t = 2.38, p = 0.023, 95% CI = 0.01, 0.06 and low FA in the genu of corpus callosum: β = 0.02, t = 2.20, p = 0.034, 95% CI = 0.00, 0.05). Whereas with high FA values, physical activity did not associate with working memory. Furthermore, we did not find evidence of white matter moderation on the link between other core executive functions and either physical activity or aerobic fitness.

**Fig 2.**
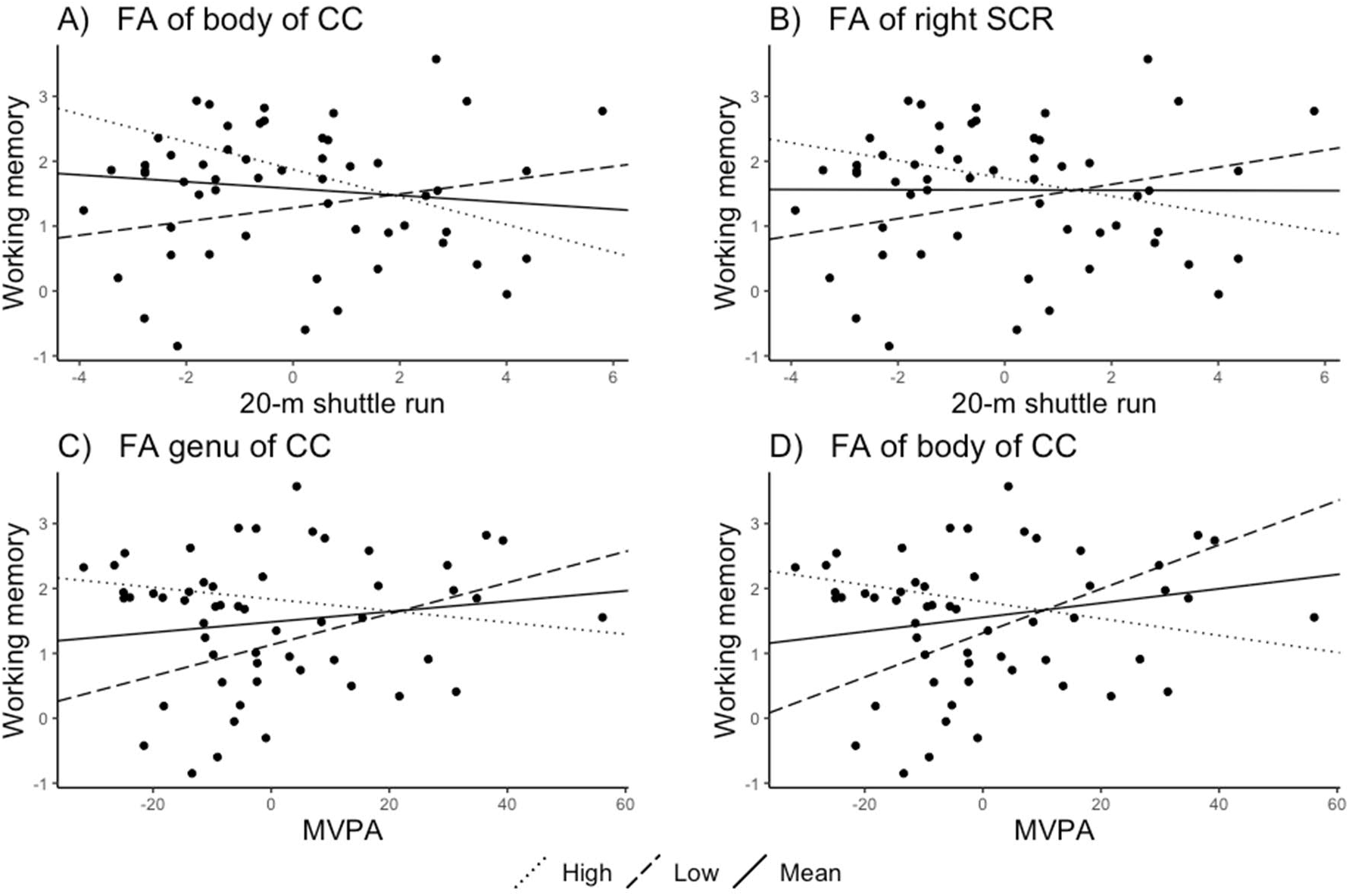
The moderating effect of FA of (High: + 1 SD, Low: −1 SD) (A) the body of corpus callosum (CC), and (B) the right superior corona radiata (SCR) regarding the relationship between working memory and the 20-m shuttle run performance. The moderating effect of FA of (C) the genu and (D) the body of corpus callosum regarding the relationship between working memory and the moderate-to-vigorous physical activity (MVPA).

## Discussion

The purpose of the current study was to determine 1) whether physical activity and aerobic fitness are related to white matter properties in adolescents and 2) whether white matter FA moderates the relationship of physical activity and aerobic fitness with core executive functions. We found that aerobic fitness (tested by a 20-m shuttle run test) is associated with white matter properties of several white matter tracts in adolescents. On the contrary, we did not find evidence for a relationship between physical activity (MVPA) and white matter properties in adolescents. Our exploratory analysis suggests that the relationship between either aerobic fitness or physical activity with working memory was moderated by FA in specific white matter tracts.

Whole-brain analysis revealed that the association between aerobic fitness and white matter FA were apparent in many white matter tracts, most robustly in the corpus callosum and the bilateral superior corona radiata. The negative associations between aerobic fitness and white matter MD and RD were even more widespread (Fig 1., Table 2). Our findings are consistent with the study conducted by Chaddock-Heyman et al. (2014) who investigated preadolescent children and demonstrated that a higher aerobic fitness was related to a greater FA in the body of corpus callosum, the superior corona radiata and the superior longitudinal fasciculus in children. The current results, however, differ from Herting et al. (2014) as they did not find evidence for a relationship between aerobic fitness and white matter FA in whole-brain analyses of 15−18-year-old male participants. The smaller number of participants in the Herting et al. (2014) study (n = 34) might not be sufficient to detect less apparent associations.

In addition to the studies of the connection between aerobic fitness and white matter FA, a few additional studies have investigated the link between physical activity and white matter in children or adolescents. Our findings that show no association between physical activity and white matter properties are in line with Herting et al. (2014) as they similarly did not find evidence of an association between MVPA and white matter FA in male adolescents. Our results are also similar to Krafft et al. (2014), in which exercise intervention did not affect white matter properties compared to the controls. In contrast to our study and the one of Herting et al. (2014), Chaddock-Heyman et al. (2018) reported an increase in white matter FA in the genu of corpus callosum as a result of an exercise intervention that did not affect aerobic fitness. However, they did not show associations between exercise and any of the other white matter tracts studied. In line with some earlier studies (Chaddock-Heyman et al., 2018, 2014; Herting et al., 2014), we did not find support for a link between physical activity and FA of the uncinate fasciculus, which was reported by Schaeffer et al. (2014). Based on the current and previous studies, there is little support for an association between physical activity and white matter in youth.

Our data suggest that fitness and physical activity have an unequal contribution to white matter structures in the brain, with associations evident only for aerobic fitness. This might be driven by genetic factors as, presumably, at least partly different genetic factors underlie fitness and physical activity. These findings also accord with our earlier observation using structural MRI, which showed that aerobic fitness, but not physical activity is related to regional grey matter volumes in adolescents (Ruotsalainen et al., 2019). It is possible that together, these results reflect partly the pronounced change that occurs in adolescent physiology during puberty. Perhaps at certain developmental time-windows, the fast changes in the brain are more strongly driven by genetic factors so that the mechanisms through which the physical activity influences the brain network are overridden. It is thus possible that the differences between the current results and some of the previously published results are due to different developmental stages of the participants.

Even though we found an association between aerobic fitness and white matter properties, we did not find a clear association between either aerobic fitness or physical activity with core executive functions. The results from previous studies that investigated these relationships differ quite extensively. For instance, aerobic fitness has been associated with executive functions (e.g. Chaddock et al., 2010; Huang et al., 2015; Westfall et al., 2018), but not all studies have shown this association (e.g. de Greeff et al., 2016; Stroth et al., 2009). So, even though two meta-analyses (Álvarez-Bueno et al., 2017; de Greeff et al., 2018) suggest that there is a small effect from exercise intervention on core executive functions, it still remains debatable (Diamond and Ling, 2019, 2016; Hillman et al., 2018).

Our results suggest that the disparity in findings may reflect underlying neurobiological factors – more precisely, the white matter integrity measured with FA. Indeed, white matter properties have been suggested to contribute to the level of improvement by cognitive interventions (de Lange et al., 2017; Engvig et al., 2012; Mackey et al., 2012) and may also represent one of the brain level targets for the physiological effects of exercise. In the current study, we did exploratory analyses to investigate whether white matter FA has an influence on the relationship between physical performance measures and cognitive function. Interestingly, moderation analysis revealed that white matter FA had a moderation effect on working memory but not on the other tested cognitive functions.

We demonstrated that at high FA levels of the body of corpus callosum and the superior corona radiata, the relationship between aerobic fitness and working memory was slightly negative and at low FA values the association was slightly positive. In other words, with high FA values, higher aerobic fitness was related to a poorer working memory performance, and with low FA values, higher aerobic fitness was related to a better working memory performance. This cross-over interaction was also seen when evaluating the interaction effect of FA of the body and genu of corpus callosum regarding their relationship between physical activity and working memory.

A more detailed simple slopes analysis, however, revealed that the influence of aerobic fitness was only significant with high FA levels and that the influence of physical activity was significant with low FA levels. This means that after reaching a certain level of FA, the fitness does not positively correlate with working memory. On the other hand, when the FA level is low, perhaps in the earlier stages of maturation in that particular white matter tract, the level of physical activity could exert a positive influence on white matter. These findings suggest that specific tracts of white matter also moderate the relationship between both aerobic fitness and physical activity with spatial working memory in adolescents. Overall, this observation will enable more precise hypotheses for future studies.

Even though the cross-sectional nature of this data does not make it possible to make causal inferences, these results raise the possibility that different levels of white matter FA at the start of the intervention may explain the individual differences in the intervention effects of the spatial working memory performance. Therefore, it is also possible that this could explain the contradictory results found in previous studies regarding exercise interventions and working memory.

## Limitations

There are a few limitations present in the current study. Firstly, the sample size might not have been large enough to detect weak associations. Secondly, even though accelerometers are widely used and provide an objective measure of physical movement, they do not consider the original fitness levels of participants. Thus, the same amount of physical activity or movement measured with accelerometers may produce different physiological responses and intensity levels for different participants. Thirdly, for the TBSS analysis, due to restrictions in this method, the averages of the imputed values were used. In addition, we assessed the level of aerobic fitness with 20-m shuttle run test, which is not a direct measure of cardiorespiratory fitness, however, it is considered to have good validity for estimating maximal oxygen uptake. We also want to highlight that the data were cross-sectional, and therefore, do not allow causal interpretations and the exploratory moderation results need to be interpreted cautiously. Finally, we studied 13–16-year-old adolescents and generalisations to other age groups may not be possible.

## Conclusions

We found that aerobic fitness and physical activity have an unequal contribution to brain white matter properties in adolescents. Aerobic fitness – assessed with a 20-m shuttle run test – positively associated with FA and negatively with MD and RD in several white matter tracts. However, we did not find these associations when studying physical activity. This result might be driven by genetic factors, which underlie fitness more strongly than those of the physical activity measures. It is also possible that only physical activity sufficient to increase aerobic fitness is needed to influence white matter. Overall, our findings concerning the exploratory moderation analysis further suggest that the level of white matter FA of specific white matter tracts may influence the relationships between either aerobic fitness or physical activity with the spatial working memory.

## Acknowledgements

This work was supported by the Academy of Finland [grant numbers 273971, 274086 and 311877] and the Alfred Kordelin Foundation. We thank Marita Kattelus, Riikka Pasanen and Jenni Silvo for their valuable help in the data collection. We also would like to thank Dr Toni Auranen and Prof. Veikko Jousmäki for providing the research infrastructure for this work.

---

**Supplementary Table 1.** Associations between the 20-m shuttle run, physical activity and core executive functions.

| Cognitive tests |  | 20-m shuttle run | MVPA |
| --- | --- | --- | --- |
| Flanker BI | $\beta$ | -17.434 | -0.972 |
|  | 95% CI | -33.937, -0.931 | -2.766, 0.823 |
|  | p-value | 0.038 | 0.282 |
|  | FDR-adjusted p-value | 0.170 | 0.483 |
|  | R<sup>2</sup> | 0.14 | 0.07 |
| Flanker BC | $\beta$ | -16.489 | -0.916 |
|  | 95% CI | -31.029, -1.949 | -2.512, 0.680 |
|  | p-value | 0.026 | 0.254 |
|  | FDR-adjusted p-value | 0.170 | 0.483 |
|  | R<sup>2</sup> | 0.14 | 0.07 |
| Flanker AI | $\beta$ | -9.973 | -0.530 |
|  | 95% CI | -20.109, 0.162 | -1.707, 0.647 |
|  | p-value | 0.053 | 0.368 |
|  | FDR-adjusted p-value | 0.170 | 0.547 |
|  | R<sup>2</sup> | 0.14 | 0.14 |
| Flanker AC | $\beta$ | -7.759 | -0.332 |
|  | 95% CI | -17.218, 1.701 | -1.415, 0.75 |
|  | p-value | 0.105 | 0.538 |
|  | FDR-adjusted p-value | 0.251 | 0.646 |
|  | R<sup>2</sup> | 0.14 | 0.14 |
| RVP | $\beta$ | 0.124 | 0.000 |
|  | 95% CI | -0.004, 0.252 | -0.016, 0.016 |
|  | p-value | 0.057 | 0.996 |
|  | FDR-adjusted p-value | 0.170 | 0.996 |
|  | R<sup>2</sup> | 0.10 | 0.04 |
| SWM | $\beta$ | 0.016 | 0.007 |
|  | 95% CI | -0.125, 0.158 | -0.009, 0.023 |
|  | p-value | 0.816 | 0.411 |
|  | FDR-adjusted p-value | 0.891 | 0.547 |
|  | R<sup>2</sup> | 0.03 | 0.04 |

N = 59 (58 for the Flanker tasks), female = 39 (38 in total for the Flanker tasks)
$\beta$ : regression coefficient, CI: confidence interval, FDR: false discovery rate, MVPA: moderate-to-vigorous intensity physical activity, BI: incompatible incongruent, BC: incompatible congruent, AI: incompatible congruent, AC: compatible congruent, RVP: rapid visual information processing, SWM: spatial working memory. The Adjusted R<sup>2</sup> denotes the adjusted proportion of the variance explained by the model.

